# Privacy-preserving integration of multiple institutional data for single-cell type identification with scPrivacy

**DOI:** 10.1101/2022.05.23.493074

**Authors:** Shaoqi Chen, Bin Duan, Chenyu Zhu, Chen Tang, Shuguang Wang, Yicheng Gao, Shaliu Fu, Lixin Fan, Qiang Yang, Qi Liu

## Abstract

The rapid accumulation of large-scale single-cell RNA-seq datasets from multiple institutions presents remarkable opportunities for automatically cell annotations through integrative analyses. However, the privacy issue has existed but being ignored, since we are limited to access and utilize all the reference datasets distributed in different institutions globally due to the prohibited data transmission across institutions by data regulation laws. To this end, we present *scPrivacy*, which is the first and generalized automatically single-cell type identification prototype to facilitate single cell annotations in a data privacy-preserving collaboration manner. We evaluated *scPrivacy* on a comprehensive set of publicly available benchmark datasets for single-cell type identification to stimulate the scenario that the reference datasets are rapidly generated and distributed in multiple institutions, while they are prohibited to be integrated directly or exposed to each other due to the data privacy regulations, demonstrating its effectiveness, time efficiency and robustness for privacy-preserving integration of multiple institutional datasets in single cell annotations.

## Introduction

Single-cell transcriptomics is indispensable for understanding cellular mechanisms of complex tissues and organisms^1–5^. As single-cell technologies rapidly developed over recent years, its experimental throughput increased substantially, allowing to profile increasingly complex and diverse samples, and accumulating vast numbers of datasets over time. Integrative analyses of such large-scale datasets originating from various samples, different platforms and different institutions globally, offer unprecedented opportunities to establish a comprehensive picture of cell landscape. To this end, various community generated large-scale atlas-level single cell reference data, such as the Human Cell Atlas (HCA^6^), Human Tumor Atlas Network^7^, BRAIN Initiative Cell Census Network^8^, Human Lung Atlas^9^, Human Gut Atlas^10^, Human BioMolecular Atlas Program (HuBMAP^11^), The Tabula Sapiens^12^, hECA^13^ etc., and recently great achievements has been made in the building of pan-tissue single-cell transcriptome atlases covering more than a million cells, including 500 cell types,across more than 30 human tissues from 68 donors^12, 14–17^.These references data facilitate the automatically cell type annotations in a supervised way without prior marker gene annotations^18–24^. It is obviously that integrating more reference datasets or combining these atlas-level data will improve the cell type annotations ^18, 19, 23, 25^, and various integration methods for single cell annotations have been presented^18, 26, 27, 28^. However, all these existing integration methods require to access the relevant reference datasets directly, which may be unavailable due to the data privacy and security issues. Currently, the privacy and political issues towards omics data transmission and sharing across different institutions or countries are gradually attracting attentions. On the one hand, countries around the world are strengthening laws to protect data privacy and security by prohibition of certain data transition across countries or organizations. Such regulations include the General Data Protection Regulation (GDPR^29^) implemented by the European Union, the Health Insurance Portability and Accountability Act (HIPPA^30^) and Health Information Technology for Economic and Clinical Health Act (HITECH^31^) enacted by U.S. and etc. On the other hand, single-cell reference datasets are generated and accumulated rapidly in different institutions around the world. It is a great demand to integrate all these institutional data globally to facilitate the establishment of the comprehensive picture of human cell reference map. However, these institutions may be required to protect data privacy and security and prohibit certain data transmission across organizations and countries by data regulation laws (Table 1). As a result, there exist an inevitable contradiction between the rapidly accumulated single cell reference data and the privacy issue among data sharing and integration. Current references integrating strategies failed to address these issues, hindered by legal restrictions on data sharing^32^. Moreover, current integrating methods ignore the problem of privacy towards single cell sequencing data, as scRNA-seq datasets of human are sensitive and likely to contain sufficient sequencing depth to call genetic variants^33^.

**Table 1.** Single cell atlases and their corresponding data regulation laws.

| Atlas | Data regulation laws |
| --- | --- |
| Human Cell Atlas<sup>6</sup> | GDPR |
| Expression Atlas<sup>36</sup> | EMBL Internal Policy for Data Protection |
| Human Tumor Atlas Network<sup>7</sup> | NIH Genomic Data Sharing Policy |
| BRAIN Initiative Cell Census Network<sup>8</sup> | NIH Genomic Data Sharing Policy |
| Human Lung Atlas<sup>9</sup> | All U.S. Federal, state and local laws and regulations |
| Human Gut Atlas<sup>10</sup> | All U.S. Federal, state and local laws and regulations |
| HuBMAP<sup>11</sup> | HuBMAP External Data Sharing Policy |
| The Tabula Sapiens<sup>12</sup> | The Tabula Sapiens Privacy Policy including GDPR |

To this end, taking the single-cell transcriptome data as an initial study, we propose *scPrivacy*, an efficient, flexible and extendable automatically single-cell type identification prototype and a proof-of-concept study to facilitate single cell annotations in a data privacy-preserving collaboration manner, by integrating multiple references single cell transcriptome data distributed in different institutions using a federated learning based deep metric learning framework. Federated learning is a collaborative paradigm in privacy-preserving computing community that enables the institutions collaboratively train a model while keeping the data in local institutions^34, 35^. We summarized existing privacy-preserving methods with their distint characteristics and explained the necessity to perform privacy-preserving computing for large scale single cell data using federated learning framework (Table 2). In addition, in our previous studies^25, 27^, metric learning was also proven to be effective for single-cell type annotation. Briefly, the basic idea of *scPrivacy* is to make each institution train their models locally and aggregate encrypted models parameters for all institutions to avoid putting raw data of all institutions together directly. We evaluated *scPrivacy* on a comprehensive set of 27 publicly available benchmark datasets for single cell type identification to stimulate the scenario that the reference datasets are rapidly generated and accumulated from multiple institutions and 15 publicly available patients datasets to simulate a large-scale real world situation that multiple hospitals collaborate together to build an automated cell type annotation system for covid-19 patients, while they are prohibited to be integrated directly or exposed to each other due to the data privacy regulations, and demonstrated its effectiveness, time efficiency and robustness for privacy-preserving integration of multiple institutional datasets.

**Table 2.** Current privacy-preserving methods and their properties.

|  | Cleartext | Software<br>Guard<br>Extensions<sup>37</sup> | Homomorphic<br>Encryption<sup>38</sup> | Secure<br>Multi-Party<br>Computation<sup>39</sup> | Federated<br>learning<sup>40</sup> |
| --- | --- | --- | --- | --- | --- |
| Risk for calling genetic variants? | Yes | No | No | No | No |
| Encrypted raw data? | No | No | Yes | Yes | No |
| Practicable for handling large scale single cell reference data? | Yes | No | No | No | Yes |
| Applicable to data transmission regulation laws? | No | No | No | No | Yes |
| Performance compared to cleartext based integration? | / | Identical | Identical | Identical | Upper bound |

## Results

### Overview of *scPrivacy*

*scPrivacy* is an efficient, flexible and extentable automatically single-cell type identification prototype to facilitate single cell annotations in a data privacy-perserving collaboration manner, by integrating multiple references single cell transcriptome data distributed in different institutions using a federated learning based deep metric learning framework. *scPrivacy* can effectively integrate information from multiple references while keeping each reference in local institutions, so as to solve the problem of data privacy protection. In particular, each institution trains its model on its local dataset and sends encrypted model parameters to server and then the server aggregates the parameters and sends back the aggregated model to institutions iteratively in training process. Specifically, *scPrivacy* comprises two main steps: model learning and cell assignment (Fig. 1 and see Methods).

**Fig. 1.**
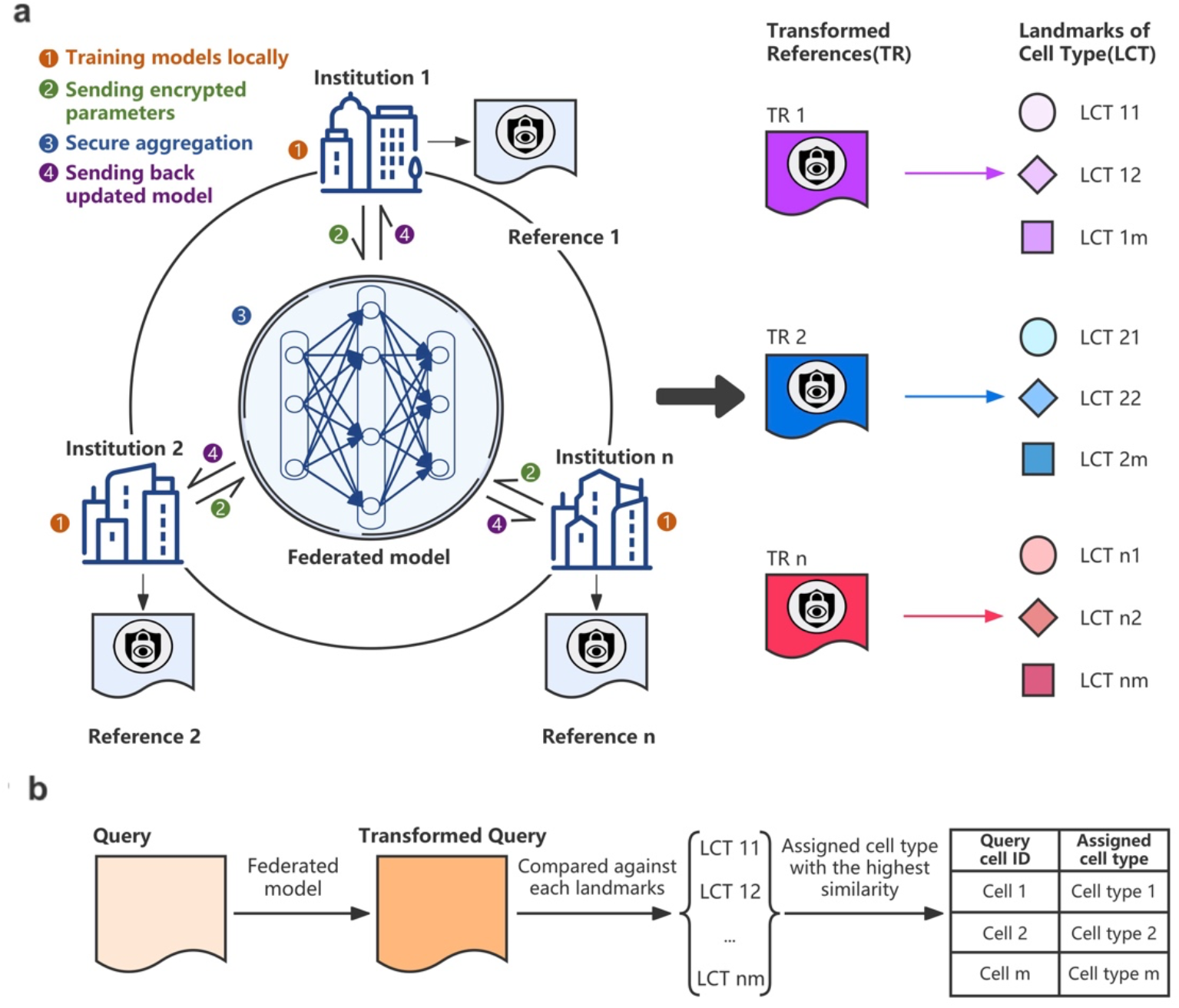
The *scPrivacy* workflow. **a**. The model learning process of *scPrivacy*. The federated model was trained with four steps: (1) Training models locally; (2) Sending encrypted parameters; (3) Secure aggregation; (4) Sending back updated model. Then cell type landmarks will be calculated for each transformed reference. **b**. The cell assignment process of *scPrivacy*. The federated model is utilized to transform the query cells. Then, the transformed query cells are compared against cell type landmarks of transformed institutional datasets, and the predicted cell type with the highest similarity among all cell type landmarks is obtained.

In the model learning stage (Fig. 1a), *scPrivacy* trains a federated deep metric learning model on multiple institutional datasets in a data privacy-preserving manner. For an individual institution, deep metric learning (DML) is applied to learn an optimal measurement fitting the relationship among cells in the reference dataset, and the N-pair loss^41^ is used as the loss function for model training. With DML, cells belong to the same type became more similar and cells belong to different types became more dissimilar. Then, *scPrivacy* extends DML to a federated learning framework by aggregating model parameters of institutions to construct an aggregated model (Fig. 1a and see Methods), which fully utilized the information contained in multiple institutional datasets to train the aggregated model while avoiding integrating datasets physically. In addition, *scPrivacy* can utilize the complementary information from different institutional datasets to boost the cell assignment performance, while also avoid the over correction of batch effect, as proven in previous study^41^. In the cell assignment stage (Fig. 1b and see Methods), the query dataset is first transformed by the federated model to the same embedding space as that of the transformed institutional datasets. Then, the transformed query dataset is assigned to proper cell types by comparing with cell type landmarks of transformed institutional datasets. Specifically, for each transformed institutional dataset, *scPrivacy* carries out a cell search by measuring the similarity between the transformed query cells and cell type landmarks of the transformed institutional datasets. Finally, the query cells are assigned to the proper cell type with the highest similarity among all landmarks of the transformed institutional datasets.

### Benchmarking *scPrivacy* with multiple institution and single institution

We firstly benchmarked *scPrivacy* with multiple institution(default), and compared it with that of *scPrivacy* with single institution to prove the benefits and necessity of integrating multiple institutional datasets. Here, “*scPrivacy* with single institution” represents that the results are achieved by training with one institution dataset and testing on another institution datasets. To this end, we collected 27 datasets (Supplementary Table S1) from four studies of three tissues to simulate the data collaboration scenario among institutions: one study on the brain, one study on the pancreas and two studies on peripheral blood mononuclear cells (PBMCs)^42, 43^. For brain tissue, the study contained four brain datasets^44, 45^ with different sources. For pancreas tissue, the study contained four commonly used pancreas datasets^46–49^. For PBMCs, the first study^43^ contained 12 datasets from 12 different sequencing platforms (“PBMC-Mereu”), and the second study^42^ contained 7 datasets from 7 different sequencing platforms (“PBMC-Ding”). In this case, each dataset of a tissue is simulated as an institution. For the strategy of *scPrivacy* with single institution, each dataset among multiple datasets was simulated as the dataset in an individual institution to train *scPrivacy* and the rest datasets were used as query datasets. For the strategy of *scPrivacy* with multiple institutions, each dataset among the multiple datasets was used as the query, and the others datasets were used to simulate distributed multiple institutional datasets and they are “virtually” integrated to train *scPrivacy*. It should be noted that in all benchmark scenarios in our study, macro-F1 score was used as the evaluation metric and only query cell types included in multiple institutional datasets were calculated. As shown in Fig. 2a and Supplementary Table S2, it can be clearly seen that *scPrivacy* with multiple institutions generally obtained great improvement compared with that of *scPrivacy* with single institution, demonstrating the importance of integrating multiple institutional datasets. To further analyze the results, we compared macro-F1 score on ‘PBMC-Mereu’ and ‘PBMC-Ding’ studies in terms of each cell type, as the two studies shared several common cell types. The results showed that *scPrivacy* achieved a better performance in almost all cell types, further demonstrating the benefits and necessity to integrate multiple institutional datasets (Fig. 2b and Supplementary Table S3).

**Figure 2.**
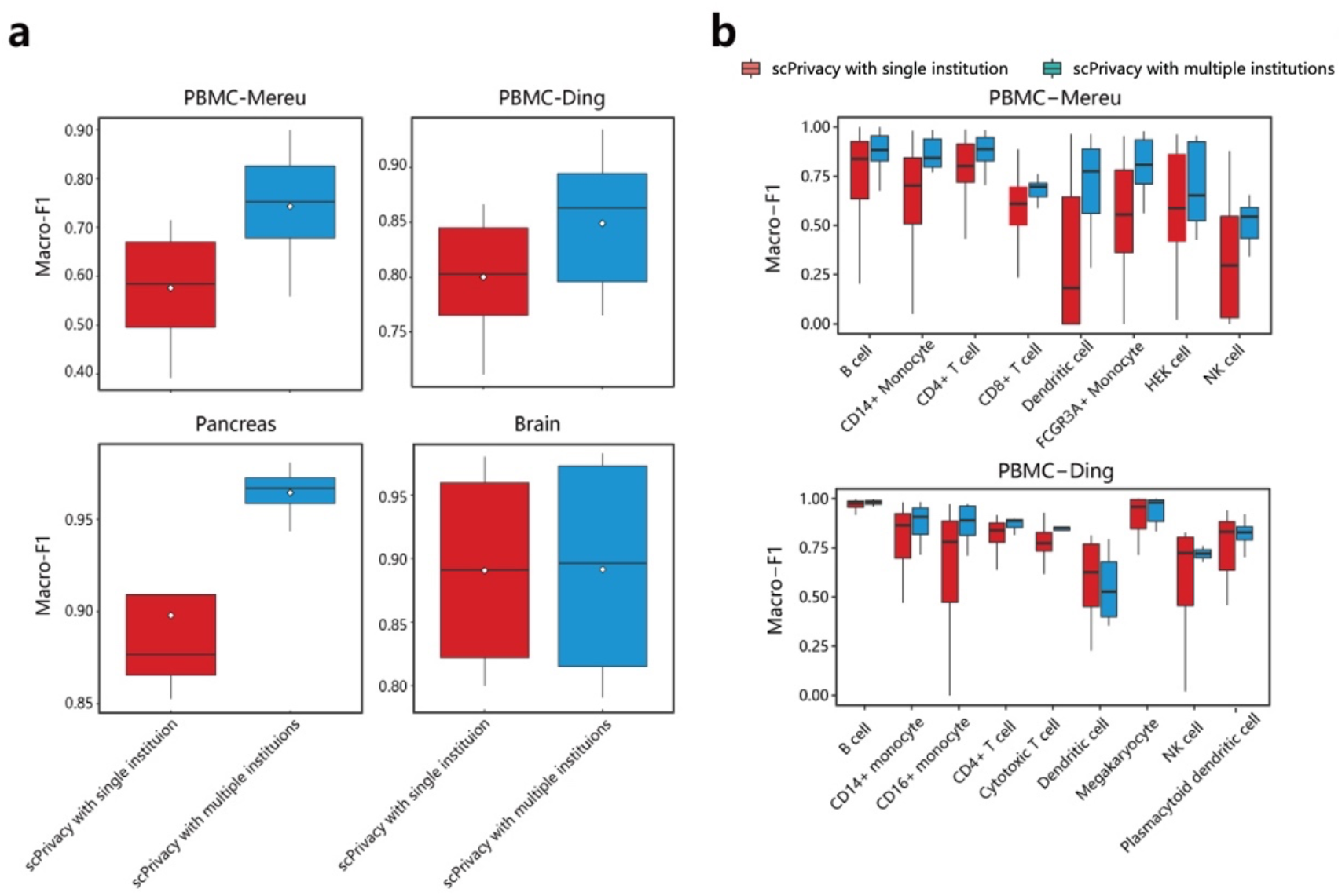
Benchmarking *scPrivacy* with single institution and multiple institutions. **a**. The macro-F1 scores of *scPrivacy* with single institution and multiple institutions for “PBMC-Mereu”, “PBMC-Ding”, “Brain” and “Pancreas” studies, respectively. The white diamond represents the mean value. **b**. The macro-F1 of each cell type for *scPrivacy* with single institution and multiple institutions on “PBMC-Ding” and “PBMC-Mereu” studies.

### Benchmarking *scPrivacy* with non-privacy-preserving multiple reference integrating methods

Then we benchmarked *scPrivacy* with existing non-privacy-preserving multi-reference based single cell type identification methods, including scmap-cluster^26^, SingleR^18^, Seurat v3^50^ and mtSC^51^. In this study, Seurat v3 applied a data-level integration strategy; scmap-cluster and SingleR applied a decision-level integration strategy, and mtSC applied an algorithm-level integration strategy (See Methods), which represent the three mainstream integrative analysis strategies for single cell assignment. The above 27 datasets in 4 studies are used as the benchmark datasets here. In this case, each dataset among the multiple datasets was treated as the query, and the others were used to simulate multiple institutional datasets. It should be noted that *scPrivacy* integrated multiple institutional datasets in a data privacy-preserving manner while other multi-reference based methods accessed all datasets to integrate them directly. The results are shown in Fig. 3a and Supplementary Table S4. We can see that *scPrivacy* achieved comparable performance to mtSC and performed better than the other methods, while it was trained in a data protection manner. Furthermore, as shown in Fig. 3b and Supplementary Table S5, we compared the macro-F1 of each cell type on ‘PBMC-Ding’ and ‘PBMC-Mereu’ studies, and achieved the consistent conclusion that *scPrivacy* achieved the comparable best performance in all four studies as those of mtSC. As a conclusion, *scPrivacy* is able to achieve a comparable best performance to integrate multiple institutional datasets, while keeping the integration in a data privacy-preserving manner.

**Figure 3.**
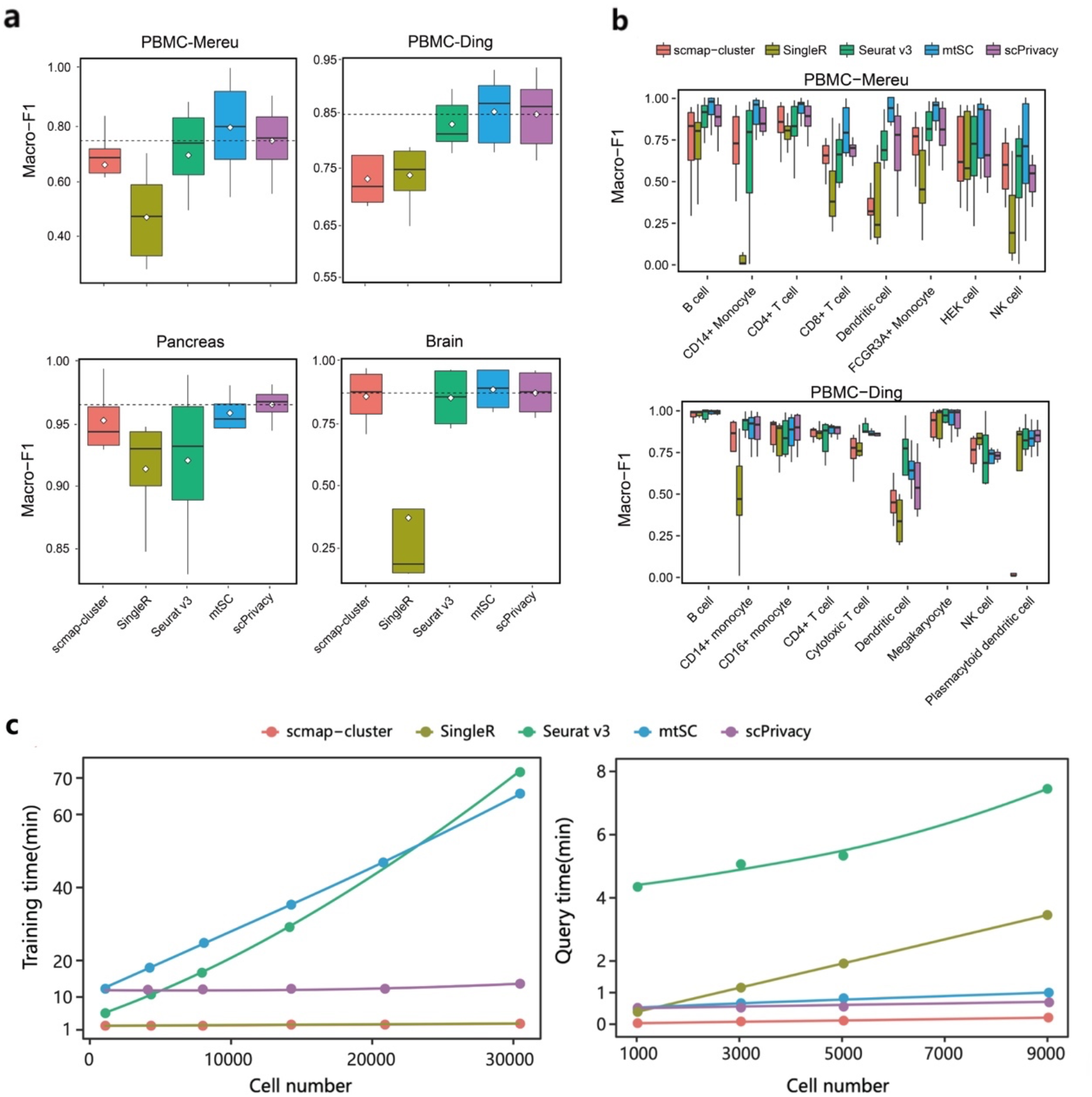
Benchmarking *scPrivacy* with non-privacy-preserving multiple reference integrating methods. **a**. The macro-F1 scores of *scPrivacy* and other existing non-privacy-preserving multiple reference integrating methods on “PBMC-Mereu”, “PBMC-Ding”, “Brain” and “Pancreas” studies, respectively. **b**. The macro-F1 of each cell type for *scPrivacy* and other existing non-privacy-preserving multiple reference integrating methods on ‘PBMC-Ding’ and ‘PBMC-Mereu’ studies. **c**. Training and query time of *scPrivacy* and other existing non-privacy-preserving multiple reference integrating methods. Solid lines are loess regression fitting (span = 2), implemented with R function geom smooth().

### *scPrivacy* consumes much less time than most multiple reference integrating methods

As the amount and size of scRNA datasets increasing rapidly, consuming time is an important concern for single cell integration and annotation. Due to the distributed properties of federated learning, the training time of *scPrivacy* only depends on the max training time among the institutions datasets while the training time of other multi-reference based methods are the sum of training time of all institution datasets. Thus, *scPrivacy* obtains a high scalability to deal with the large-scale data integration and model training, which serves a very challenging issue in the building of large-scale atlas level single cell reference data. The following training time and query time comparisons further proved this point (Fig. 3c, Supplementary Table S6 and S7): (1) *scPrivacy* consumes much less time than Seurat v3 and mtSC in the training process. More importantly, the time consumed by *scPrivacy* does not increase exponentially or linearly, since it only depends on the max training time among institutions datasets, indicating its potential ability to handle large-scale datasets. (2) For querying process, *scPrivac*y is also very fast (<1 min for 9000 query cells) and consumes much less time than all the other methods except for scmap-cluster, since scmap-cluster is the simplest method while expense of accuracy substantially.

### Robustness validation of *scPrivacy*

We then validated the robustness of *scPrivacy*. We firstly investigated the impact of the number of institutions on the performance of *scPrivacy*. We considered each dataset in “PBMC-Ding” as an institution dataset to simulate the scenario. In this test, *scPrivacy* was trained with different numbers of institution datasets, and then the corresponding macro-F1 score was calculated to show the trend of the performance as the number of institution datasets increased. Specifically, each time, we randomly selected one dataset from the 7 datasets as the query dataset and selected 1 to 6 datasets without replacement from the remaining datasets as the institutional reference datasets. This process was repeated 5 times to reduce randomness. As shown in Fig. 4a and Supplementary Table S13, we can see that *scPrivacy* generally performs increasingly better as the number of institutions increases. The macro-F1 scores for each dataset as query dataset can be found in Supplementary Figure S1. Specifically, if the performance is relatively lower in the beginning, the improvement becomes more evident as the number of institution datasets increases. Such an improvement obtained by increasing institution datasets is very important for developing an effect and robust cell annotation system in the era of explosive growth of single-cell datasets. Together with the fast training of *scPrivacy*, it is expected to integrate a growing number of institution datasets to obtain an increasingly better cell annotation in a privacy-preserving manner.

**Figure 4.**
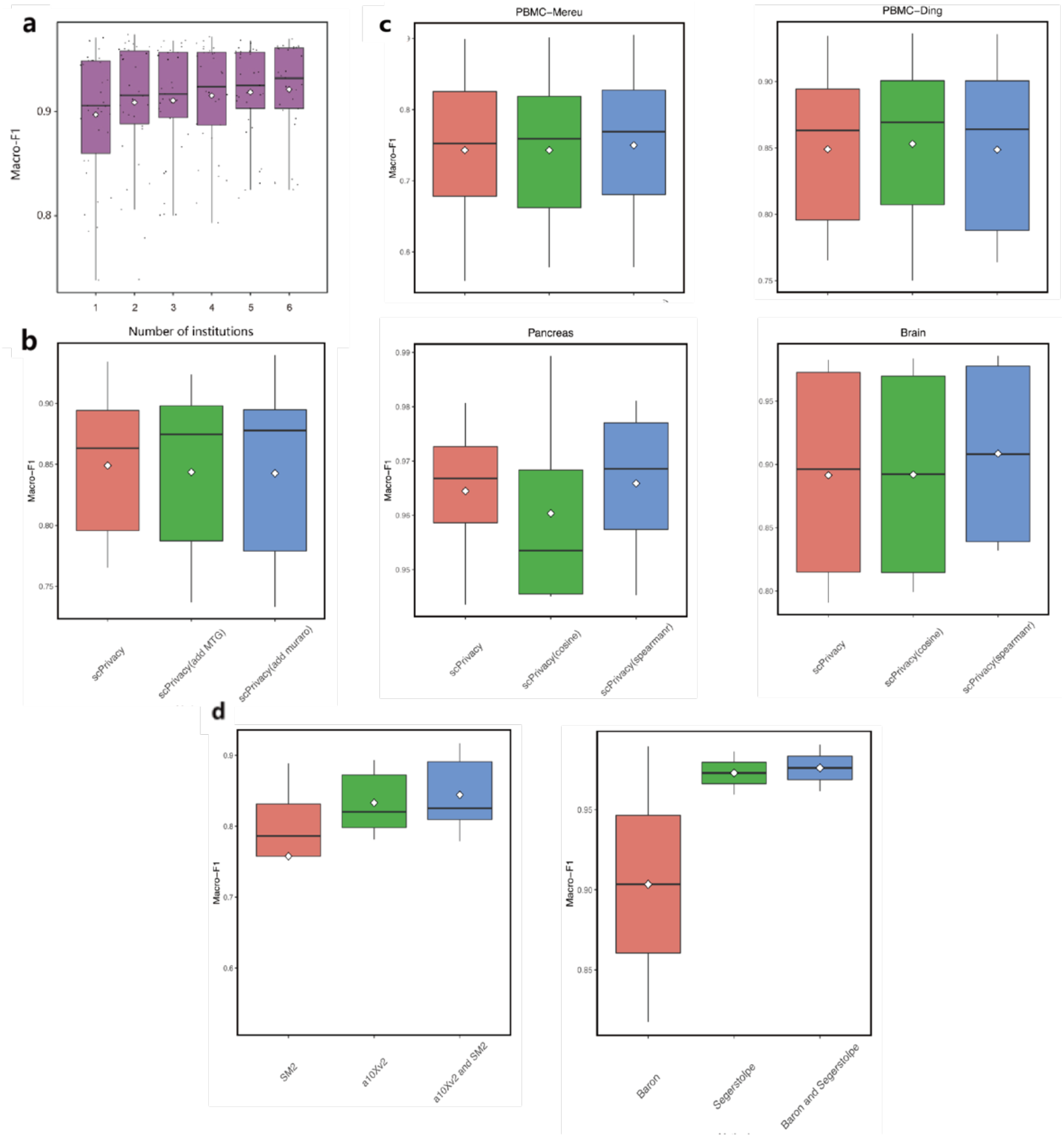
Robustness validation of scPrivacy. **a**. The performance of *scPrivacy* as the number of institution datasets increases. We considered each dataset in “PBMC-Ding” as an institution dataset to simulate the scenario, and showed the macro-F1 scores for overall datasets. **b**. The macro-F1 scores of simulation that the data of an institution is from a different tissue while the others are from the same tissue. Each dataset among PBMC-Ding was treated as the query, and the others and a different tissue dataset (the brain dataset MTG or pancreas dataset Muraro) were used to simulate multiple institutional datasets. **c**. The macro-F1 scores of *scPrivacy* using different similarity metrics including Pearson correlation coefficient, cosine similarity and Spearman correlation coefficient. **d**. The macro-F1 scores of simulation that the data volumn of institutions vary greatly. In PBMC-Ding datasets, dataset a10Xv2 and SM2 were considered as the institutions datasets and the others were treated as the query. In pancreas datasets, dataset Baron and Segerstolpe were considered as the institutions datasets and the others were treated as the query. The white diamond represents the mean value.

Then we explored the influence of similarity metrics for query cell assignment. Another two common similarity metrics cosine similarity and spearman correlation coefficient were tested here. As shown in Fig. 4c, using different similarity metrics has little influence on the results and *scPrivacy* is robust to different similarity metrics. In addition, we explored the influence of the data of an institution completely different from that of other institutions. In our understanding, different tissue datasets are heterogeneous and different from each other. To explore the scenario that the data of one institution is completely different from that of other institutions, it is a proper choice to simulate the data of an institution is from a different tissue while the others are from the same tissue. To this end, each dataset among PBMC-Ding was treated as the query, and the others and a different tissue dataset (the brain dataset MTG or pancreas dataset Muraro) were used to simulate multiple institutional datasets. As shown in Fig. 4b, when the data of one institution is completely different from that of other institutions, the model performance will only decrease slightly.

Finally, we explored the influence of greatly varying data volume across institutions. To simulate the scenario, we selected datasets whose data volumes vary greatly as the institution datasets. In PBMC-Ding datasets, dataset a10Xv2 (9683 cells) and SM2 (475 cells) were considered as the institutions datasets and the others were treated as the query. In pancreas datasets, dataset Baron (8562 cells) and Segerstolpe (2126 cells) were considered as the institutions datasets and the others were treated as the query. As shown in Fig. 4d, in general, *scPrivacy* with multiple institutions generally obtained improvement compared with that of *scPrivacy* with single institution, which is consistent with the conclusion in Fig. 2. In addition, the results showed that the improvement of the institution with the less data volume is larger than that of the institution with the larger data volume. Taken together, *scPrivacy* is robust to the number of institutions, similarity metrics, data heterogeneity and data volumn, further indicating that it has the ability to handle the complex situation in real world.

### A large-scale simulation of collaborations between multiple hospitals for covid-19 patient cell annations with *scPrivacy*

In this study, we made a large-scale simulation of a real world scenario that multiple hospitals collaborate together to build an automated cell type annotation system for covid-19 patients (Fig. 5a). It is obvious that the patient data always has patient privacy issue and hospitals can not share patient data with other institutions if they are not approved by patients. Zhang et al^52^. constructed a large-scale PBMC single cell transcriptome atlas which consists of 196 individuals in 5 disease stages from 39 institutes or hospitals. As several integrating methods (such as Seurat v3) can not handle the simulation using all individuals, we randomly selected 15 large-scale individuals datasets (> 6000 cells) satisfying certain criteria (see Methods) (Supplementary Table S12) from different hospitals. The similar benchmark strategies descripted aforementioned were also adopted here. We firstly benchmarked *scPrivacy* with multiple hospitals with that of *scPrivacy* with single hospital. As shown in Fig. 5b, c and Supplementary Table S7, S8, *scPrivacy* with multiple hospitals also obtained great improvement in general, especially in terms of cell types which are more difficult to be distinguished compared with that of *scPrivacy* with single hospital, demonstrating the effectiveness of integrating multiple hospital patients datasets. Then we benchmarked *scPrivacy* with existing non-privacy-preserving data integration methods. As shown in Fig. 5d, e and Supplementary Table S9, S10, we can see that *scPrivacy* achieved state-of-the-art performance in all cell types while it was trained in a data protection manner. As a conclusion, this large-scale real world application simiultion also proved that *scPrivacy* is able to handle large-scale data integration issue and achieve state-of-the-art performance to integrate multiple institutional datasets, while keeping the integration in a data privacy-preserving manner.

**Figure 5.**
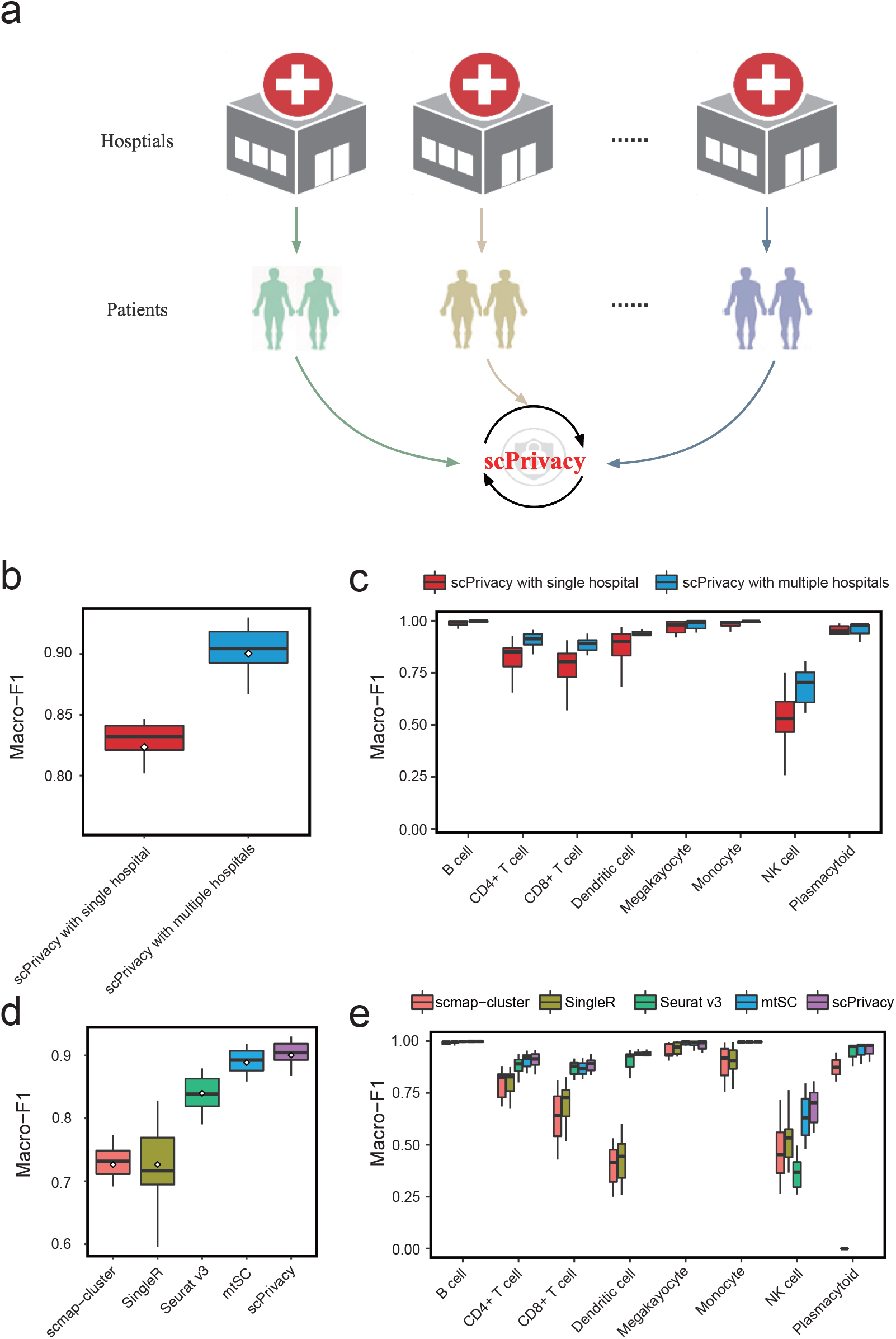
A large-scale real world simulation that multiple hospitals collaborate together to build an automated cell type annotation system for covid-19 patients with *scPrivacy*. **a**. The workflow of the simulation. **b**. The macro-F1 scores of *scPrivacy* with single hospital and multiple hospitals. **c**. The macro-F1 of each cell type for *scPrivacy* with single hospital and multiple hospitals. **d**. The macro-F1 scores of *scPrivacy* and other existing non-privacy-preserving multiple reference integrating methods. **e**. The macro-F1 of each cell type for *scPrivacy* and other existing non-privacy-preserving multiple reference integrating methods. The white diamond represents the mean value.

## Discussion

In this study, we present the first and generalized federated deep metric learning-based single cell type identification prototype *scPrivacy* to facilitate single cell annotations by integrating multiple institutional datasets in a privacy-preserving manner. As scRNA-seq datasets grow exponentially, multiple institutional datasets can be integrated to build a more comprehensive, effective and robust cell annotation system. Traditional multi-reference based methods faced the problem of data privacy protection. *scPrivacy* solves this issue by federated learning. Specifically, *scPrivacy* trains each institution dataset locally and aggregates encrypted model parameters for all institutions instead of putting raw data of all institutions together to train a model. We evaluated *scPrivacy* on a comprehensive set of publicly available benchmark datasets for single cell type identification to stimulate the scenario that the reference datasets are rapidly generated and distributed in multiple institutions, while they are prohibited to be integrated directly or exposed to each other due to the data privacy regulations, and demonstrated its effectiveness for privacy-perserving integration of multiple institutional datasets. A large-scale real world simulation that multiple hospitals collaborate together to build an automated cell type annotation system for covid-19 patients with *scPrivacy* was also demonstrated. Collectively, *scPrivacy* is time efficient, performing increasingly better as the number of institution datasets increases, and robust to the number of institutions, similarity metrics, data heterogeneity and data volumn, which is of great potential utility in various real world applications to build single cell atalas in a privacy-preserving way.

In addition to scRNA-seq technology, other single cell omics, such as scATAC-seq and etc., are developing rapidly and vast numbers of datasets will accumulate in the future. Integrating these multi-omics datasets has the potential to provide a more comprehensive picture of basic biological processes. However, privacy issue is still an unavoidable problem for data sharing and integration. *scPrivacy* can be easily extended to other single cell omics for privacy-preserving integration. In addition, the privacy-preserving computing technologies are evolved rapidly, although federated learning is a suitable solution for large-scale privacy-preserving data integration, it still needs a central server to aggregate model parameters from clients. Recently, blockchain-based federated learning, such as swarm learning^53, 54^, is developed which doesn’t need the central server, however, its efficiency is waiting to be further evaluated. Nevertheless, a privacy-preserving integration of multiple institutional data globally to build a more comprehensive cell landscape is a challenging issue that must be faced under the global data sharing protection and regulations. We therefore call for attention to such problems and efficient privacy-preserving system for cell annotations are needed to be carefully designed.

## Methods

### Benchmark data collection

We evaluated *scPrivacy* on 42 reference based single-cell type identification benchmark datasets which were curated from five studies including three tissues: peripheral blood mononuclear cells (PBMCs)^42, 43, 52^, the brain^44, 45^ and the pancreas^46–49^ (Supplementary Table S1 and S12). For all datasets, cell types with less than 10 cells were removed since they do not contain enough information and are unreliable for subsequent assignment. Cells labeled “alpha.contaminated”, “beta.contaminated”, “gamma.contaminated” and “delta.contaminated” in the dataset generated from Xin et al.^49^ were removed because they likely corresponded to cells of lower quality. Cells labeled “not applicable” in the dataset generated by Segerstolpe et al.^48^ were removed. “L6b”, “Pvalb”, “Sst” and “Vip” cell types in the dataset generated from Tasic et al.^45^ were retained to match the names of cell types in the other three brain datasets^44^. For “PBMC-Mereu”^43^, 12 datasets were used and Smart-seq2-based dataset was excluded, which was too small (Supplementary Table S1).

### Data preprocessing

The data preprocessing step of *scPrivacy* consists of three parts: cell quality control, rare cell type filtering and gene expression profile formatting. *scPrivacy* evaluates the cell quality on strict criteria. In particular, the number of genes detected requires >500, the number of unique molecular identifiers induced requires >1500, and the percentage of mitochondrial genes detected requires <10% among all genes. Only cells satisfying all three criteria are reserved. The quality control of Zhang’s study datasets were processed with the quality control procedures in their paper^52^. Then, all the datasets were normalized by scaling to 10,000 and then with log(counts+1). Finally, all the datasets are processed into an identical format, i.e., expression profiles with the union of the genes in all institution datasets. The column of the gene in query dataset will be filled with zeros if the gene is not in the gene union of the institution datasets.

### The selection of the simulation datasets for collaboration among multiple hospitals in the cell annotations of covid-19 patient

We filtered the datasets not statisfying following criteria. In particular, the number of cells requires > 6000 to gurantee the large scale of dataset, the number of cell types requires > 6 to gurantee the quality of dataset. We randomly selected 15 individuals datasets statisfying criteria in various disease stages from different hospitals and made sure that there are 3 individuals datasets in each disease stage. The details of selected datasets can be found in Supplementary Table S12.

### Model learning of *scPrivacy*

In the model learning stage, *scPrivacy* utilizes federated deep metric learning algorithms to train the federated model on multiple institutional datasets in data privacy protection manner. We remove batch effects for datasets by sending the gradients learned from each dataset, then the aggregated model can utilize the information of the same cell type from different datasets so as to uncover the common biological information and remove the batch effect, which is similar to the idea of *mtSC* to remove batch effect^25^. For a single institution, we use deep metric learning (DML) as the training algorithm and N-pair loss^41^ is used as the loss function. DML is applied to learn an optimal measurement fitting the relationship among cells in the reference dataset. With the measurement learned from DML, cells with the same label become more similar and cells with different labels become more dissimilar. The DML neural network for a single institution contains an input layer, a hidden layer and an output layer. The nodes of input layer equal to the genes of the reference while the nodes of the hidden layer and output layer are 500 and 20, respectively.

The application of the *N*-pair loss consists of two parts: batch construction and calculation. For the batch construction of the *N*-pair loss, 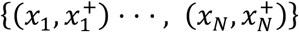 is defined as *N* pairs of cells from *N* different cell types, in which *x*_*i*_≠*x*_*j*_ ∀ *i*≠*j*. Then, *N* tuples denoted by 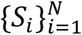 are built from the *N* pairs, where 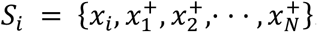. Here, *x*_*i*_ is the query for 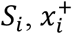 is a positive example, and 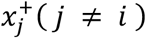 are the negative examples. *x*_*i*_ and 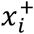 are two cells of the same cell type, and 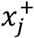 are the cells with different cell types different from *x*_*i*_.

The calculation of the *N*-pair loss can be formulated as follows:

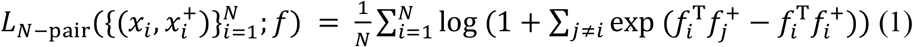

in which *f*(·; *θ*) is an embedding kernel defined by a deep neural network, *f*_*i*_ and 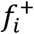 are embedding vectors of two cells of the same cell type and 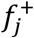 are embedding vectors of cells whose cell types are different from *x*_*i*_.

*scPrivacy* extends DML to a federated learning framework. Define *N* institutions {*F*_1_, …*F*_*N*_} and their respective data {*D*_1_, …*D*_*N*_}. A federated learning framework allows institutions to collaboratively train a model and no institution *F*_*i*_ expose its data *D*_*i*_ to others.

As shown in Fig. 1, in *scPrivacy*, institutions datasets learn a federated model collaboratively with the help of a server. To train a federated model, our training process can be divided into the following four steps^40^:

- Step 1: Each institution trains its model on its own dataset with DML;
- Step 2: Institutions encrypt their own model parameters and send encrypted parameters to server;
- Step 3: Server performs secure aggregation for encrypted parameters;
- Step 4: Server sends back the aggregated model parameters to institutions and institutions update their models with the decrypted aggregated model parameters.

In our implementation of *scPrivacy*, we used the Crypten, a ML framework built on PyTorch^55^ to implement encryption of model parameters and their operations in secure aggregation with secure multi-party computation^39^ which is adopted as an encryption technique here. At the central server, we adapted widely-used FedAvg^56^ algorithm to aggregate model parameters. The basic idea of the algorithm to aggregate model parameters is to average model parameters (weight parameters *w* and bias parameters *b*) of the neural network models for institutions. Institutions update their models by replacing model parameters with the decrypted aggregated model parameters.

### Model parameters of *scPrivacy*

The neural network model was implemented with PyTorch. We use Adam optimizer to update parameters of the network via backpropagation. The learning rate was set to 0.0005. The number of training epochs was set to 300. The L2 regularization rate was set to 0.05.

### Query cells assignment

First, the query dataset was scaled to 10,000 and normalized with log(counts+1). The column of the gene will be filled with zeros if the gene is not in the gene union of the institution datasets. Next, the query cells were transformed by the federated model to the same embedding space as that of the transformed institution datasets. Then, the transformed query dataset was assigned to proper cell types by comparing with cell type landmarks of transformed institution datasets. Specifically, for each transformed institution dataset, *scPrivacy* carried out a cell search by measuring similarity between transformed query cells and cell type landmarks of the transformed institution datasets. Pearson correlation coefficient was adopted to calculate similarity as in our previous study^22^. Finally, the query cells obtained the predicted cell types with the highest similarity among all landmarks of the transformed institution datasets.

### Benchmarking existing tools

To evaluate the performance of *scPrivacy*, four classical non-privacy-preserving data-integration methods were compared: Seurat v3^23^, scmap-cluster^19^, SingleR^18^ and mtSC^25^. Seurat v3 applied a data-level integration strategy, where multiple datasets are integrated into one dataset directly; scmap-cluster and SingleR applied a decision-level integration method, where the final assignment results are ensembled by individual assignment results; and mtSC applied an algorithm-level integration strategy, where efficient algorithms are designed for model integration, while keeping datasets separately. However, all these methods are needed to access the individual institutional data directly. To be fair, all these methods were allowed to access all institutional datasets directly while *scPrivacy* integrated multiple institutional datasets in a data privacy-preserving manner. Each dataset among the multiple datasets in a study was treated as the query, and the others were used to simulate multiple institutional datasets. For scmap-cluster, ‘threshold = 0’ was set to not assign query cells to ‘unassigned’. For SingleR, as the fine-tuning process of SingleR is extremely time consuming, ‘fine.tune = FALSE’ was set. For Seurat v3, all parameters were the defaults. For mtSC, all parameters were the defaults. In all tests, Seurat v3, scmap-cluster and SingleR and were trained and tested with CPU Intel Xeon Platinum 8165 2.3-3.7GHz. The deep learning based methods mtSC and *scPrivacy* were trained with GPU 1080Ti and tested with the same CPU as the other methods. To further prove the effectiveness of *scPrivacy*, we also benchmarked *scPrivacy* with the decision-level strategy (Supplementary Figure S2).

### Evaluation criteria

The macro-F1 score was used as evaluation criteria. Despite integration tasks, we calculated the macro-F1 score for each query dataset just like the calculation for no-integration tasks. First, the precision and recall of each cell type were calculated. Then, the macro-F1 score was calculated as listed below:

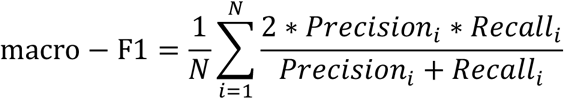

in which *N* denotes the number of cell types in a dataset and *Precision*_*i*_ and *Recall*_*I*_ are the precision and recall of the *i*-th cell type in the dataset.

## Data availability

The 42 reference based single-cell type identification benchmark datasets were curated from five studies including three tissues: peripheral blood mononuclear cells (PBMCs)^42, 43, 52^, the brain^44, 45^ and the pancreas^46–49^ (Supplementary Table S1 and S12). The four pancreas datasets^46–49^ and one of the brain datasets^45^ were collected in previous work of scmap^19^ (https://hemberg-lab.github.io/scRNA.seq.datasets), and the other three brain datasets and seven datasets in “PBMC-Ding”^42^ were curated from the following benchmark study^57^ (https://doi.org/10.5281/zenodo.3357167). The 12 datasets in “PBMC-Mereu”^43^ were collected from GSE133549, and the corresponding RData file can be downloaded in https://www.dropbox.com/s/i8mwmyymchx8mn8/sce.all_classified.technologies.RData?dl=0. All these datasets were converted into Bioconductor SingleCellExperiment (http://bioconductor.org/packages/SingleCellExperiment)class objects. The 15 datasets of Covid-19 patients were collected from GSE158055 (Supplementary Table S12).

## Code availability

*scPrivacy* is developed as a python package for simulations, which is available at https://github.com/bm2-lab/scPrivacy.

## Acknowledgements

This work was supported by the National Key Research and Development Program of China (2021YFF1200900 and 2021YFF1201200), National Natural Science Foundation of China (31970638 and 61572361), Shanghai Artificial Intelligence Technology Standard Project (19DZ2200900), Shanghai Shuguang Scholars Project, WeBank Scholars Project, and Fundamental Research Funds for the Central Universities.

## Author contributions

Q.L, Q.Y. and L.X.F. proposed the idea. S.Q.C implemented models and analyzed data. B.D. and C.T curated the benchmark datasets. B.D, S.G.W, Y.C.G and S.L.F contributed by visualized and analyzed results. Q.L, S.Q.C and B.D wrote the manuscript with assistance from other authors.

